# Neuroinflammation underlies the development of social stress induced cognitive deficit in sickle cell disease

**DOI:** 10.1101/2024.01.24.577074

**Authors:** S’Dravious A. DeVeaux, Sofiya Vyshnya, Katherine Propsom, Oluwabukola T. Gbotosho, Asem S. Singh, Robert Z. Horning, Mihika Sharma, Anil G. Jegga, Liang Niu, Edward A. Botchwey, Hyacinth I. Hyacinth

## Abstract

Cognitive deficit is a debilitating complication of SCD with multifactorial pathobiology. Here we show that neuroinflammation and dysregulation in lipidomics and transcriptomics profiles are major underlying mechanisms of social stress-induced cognitive deficit in SCD. Townes sickle cell (SS) mice and controls (AA) were exposed to social stress using the repeat social defeat (RSD) paradigm concurrently with or without treatment with minocycline. Mice were tested for cognitive deficit using novel object recognition (NOR) and fear conditioning (FC) tests. SS mice exposed to RSD without treatment had worse performance on cognitive tests compared to SS mice exposed to RSD with treatment or to AA controls, irrespective of their RSD or treatment disposition. Additionally, compared to SS mice exposed to RSD with treatment, SS mice exposed to RSD without treatment had significantly more cellular evidence of neuroinflammation coupled with a significant shift in the differentiation of neural progenitor cells towards astrogliogenesis. Additionally, brain tissue from SS mice exposed to RSD was significantly enriched for genes associated with blood-brain barrier dysfunction, neuron excitotoxicity, inflammation, and significant dysregulation in sphingolipids important to neuronal cell processes. We demonstrate in this study that neuroinflammation and lipid dysregulation are potential underlying mechanisms of social stress-related cognitive deficit in SS mice.

**Key Points:**

1. Neuroinflammation and lipid dysfunction are potential underlying mechanisms of social stress-related cognitive deficit in SCD patients.
2. Mitigating or ameliorating the impact of cognitive deficits in SCD needs to consider the biological changes already created by exposure to social stress.

**Novelty of our Findings:** We show for the first time, that neuroinflammation along with changes in the brain lipidome and transcriptome, are underlying biological mechanism contributing to the development and potentially progression of cognitive impairment among sickle cell patients. These findings also provide for the first time, a mechanistic basis for an earlier reported observation of a higher likelihood of having lower intelligence quotient scores among children with sickle cell disease exposed to social stress in the form of low parental socioeconomic status.

## Introduction

Sickle cell disease (SCD) is a common inherited blood disorder that affects approximately 100,000 Americans.^1^ SCD is caused by a point mutation in the gene for the β-globin subunit of hemoglobin. This mutation causes the hemoglobin to polymerize in conditions of low oxygen tension, causing the red blood cells (RBCs) to assume a sickle morphology.^2, 3^ Sickle RBCs are more fragile and prone to hemolysis - leading to anemia, the resulting free heme also initiates and propagates an inflammatory cascade that lead to vaso-occlusion,^2, 3^ and end organ damage.^4^

The cerebrovascular effects of SCD include stroke, cerebral macro and micro vascular abnormalities, and silent cerebral infarctions (SCIs).^5, 6^ Strokes and SCIs have been linked to cognitive impairment in SCD. However, recent studies have found cognitive dysfunction in children^7–9^ and adults^10–12^ even in the absence of MRI-detectable cerebral injury.^7, 8, 10^ Children with SCD typically have lower full-scale IQ scores, poorer academic achievement, and impaired processing speed.^7, 8^ Similarly, adults with SCD exhibit impairments in processing speed, working memory, global cognitive function, and executive function.

The mechanism underlying cognitive impairment in SCD is not well understood and one possibility is that individuals with SCD are hypersensitive to social stressors (to which individuals with SCD are exposed) which interact with biological factors leading to the development of cognitive deficit. Individuals with SCD often belong to lower socioeconomic classes with associated lower family educational attainment and income. The impact of social stress on cognitive function in SCD was recently demonstrated by several studies.^6, 13–15^ In a study by King *et al.* they reported that social stressors in the form of lower parental education levels and lower family income – had a similar albeit slightly more severe impact on cognitive function compared to biological factors – such as the presence of SCI, anemia, and age.^8, 9^ Studies in the general human population and in non-sickle cell mouse models, have shown a link between social stress and neuroinflammation. The functional effects of neuroinflammation on the brain include the development of cognitive impairment as well as neuropsychological abnormalities, such as anxiety and depression.^16^ as well as learning and memory impairments ^17–19^. Hence, neuroinflammation may be a possible mechanism for stress-induced cognitive abnormalities in SCD.

Neuroinflammation is also mediated by multiple factors, including sphingolipids and genetics. Sphingolipids are a class of bioactive lipids that participate in cell signaling. In the brain, sphingolipids modulate cytokine release and astroglia activation.^20^ Studies have shown that imbalances in the sphingolipid metabolism and distribution of lipids in the brain are associated with impaired memory and learning in both humans and animal models.^21–26^ Furthermore, enzymes in the sphingolipid pathway – such as sphingosine kinases, sphingosine-1-phosphate lyase, and sphingomyelinases – are involved in synaptic communication, learning, and memory, as well as in the regulation of other enzymes (e.g., COX2) that synthesize both pro-inflammatory and anti-inflammatory lipid mediators. Changes in the enzymatic activity have been implicated in the development of neuroinflammation and neurodegenerative diseases, such as Alzheimer’s and amyotrophic lateral sclerosis.^27^ ^28^

We have previously shown that aging and neuroinflammation contributes to cognitive impairment in SCD mouse model.^29^ In the present study, we demonstrate the role of sociological stress in cognitive and neurobehavioral deficit in SCD and show that neuroinflammation is a likely underlying mechanism. We used the repeat social defeat (RSD) paradigm, as a social stress model, as previously described.^17^ Our overall hypothesis is that *stress-related cognitive impairment in sickle cell disease is mediated by neuroinflammation and inimical changes in the brain lipidomic and transcriptomic profiles compared to controls.* Furthermore, that treatment with minocycline, an “anti-neuroinflammatory” drug, during RSD exposure in mice will reduce neuroinflammation and improve cognitive and behavioral function.

## Methods

### Study design and overall methods

This study aims to examine the mechanism underlying the development of cognitive deficit in SCD with exposure to social stress, by using the repeat social defeat (RSD) paradigm in sickle cell mice. Repeat social defeat was carried out by introducing a male intruder mouse (an aggressor) into an established cage of 3, 6 months old male sickle cell (N = 10) or control (N = 10) mice, every day, for two hours (5 – 7 PM), for six consecutive days. Age and sex-matched control (sickle cell (N = 10) and non-sickle cell (N = 10) mice) cages were set up but without aggressor mice. On the seventh day, mice were tested for cognitive/behavioral deficit using novel object recognition (NOR) and fear conditioning (FC) test paradigms. Except for the aggressors, all mice used were Townes humanized sickle cell (SS) and control (AA) mice.

To test the hypothesis that neuroinflammation is an underlying mechanism, a second cohort of SS (N = 35) and AA (N = 32) mice were randomly assigned (half from each genotype group) to receive oral (administered in drinking water) minocycline treatment (90mg/kg) or placebo (plain drinking water). Mice within each treatment arms were randomly assigned to RSD exposure or no RSD exposure. Minocycline treatment was started one day prior to the day of commencing RSD and co-terminated on the same day as the final RSD session. The minocycline dose was kept constant by adjusting the amount administered daily, using the water/drug consumption from the previous day. Cognitive/behavioral testing was performed as before, starting the next day after day 6 of RSD and day 7 of treatment.

In both experiments, the mice were randomized to histological analysis or molecular (bulk RNA sequencing and lipidomics) studies and sacrificed 1-2 days after completion of behavioral testing and brains were extracted for the assigned analysis. Cellular evidence of neuroinflammation in the hippocampus/dentate gyrus was determined using immunohistochemistry to quantify peripheral immune cell infiltrates (CD45^+^ (bone marrow-derived microglia), CD3^+^ (T-cell density), B220^+^ (B-cell density), and Iba1^+^ (activated microglia).

### Animals

We used the Townes sickle cell mouse model, which included humanized sickle cell mice (SS or HbSS) and humanized control mice (AA or HbAA).^30^ Mice were provided food and water ad libitum, housed in a 12-hour light and dark cycle, and their health statuses were monitored closely throughout the study. All experiments were approved by the Institutional Animal Use Committee at Emory University and the University of Cincinnati.

A more detailed description of the study methods is in the methods section of the Online Supplementary material. And data reporting is in accordance with the ARRIVE guidelines.

## Results

Experimental groups in this study are defined as follows in the results section: humanized control mice (AA) and Townes sickle (SS) mice, and also denote animals not exposed to stress or treated with minocycline; AA+RSD and SS+RSD denote mice exposed to stress (RSD); AA+minocycline and SS+minocycline denote mice not exposed to RSD but treated with minocycline; AA+RSD+minocycline and SS+RSD+minocycline denotes mice treated with minocycline 1 day prior to and during exposure to stress.

Figure 1 shows comparison of the groups on measure of anxiety (open-field test) and cognitive function (percent preference or freezing). In Fig. 1A**-B**, overall, we see that SS mice that were not exposed to RSD or drug treatment showed more evidence of anxiety compared to AA mice that were not exposed to RSD, indicated by the shorter distance traveled (Fig. 1A day 1: 19,116mm in AA vs. 12,593mm in SS), and relatively shorter time spent in the middle of the open field (Fig. 1B, day 1: 33.94s in AA vs. 24.06s in SS). Furthermore, SS mice exposed to RSD showed more evidence of anxiety compared to AA mice exposed to RSD (distance traveled: day 1: 12,593mm in SS vs. 9,274 cm in SS+RSD; time in the middle of the open field: day 1: 24.1s in SS vs. 32.3s in SS+RSD) or to SS or AA mice. SS+RSD+minocycline mice showed abrogation of the development of anxiety (distance traveled: day 1: 13,276mm in SS+RSD+Minocycline vs. 9,274mm in SS+RSD; time in open field: day 1: 39.6s in SS+RSD+Minocycline vs. 32.28s in SS+RSD: Figs. 1A and B) compared to untreated SS+RSD mice.

**Figure 1.**
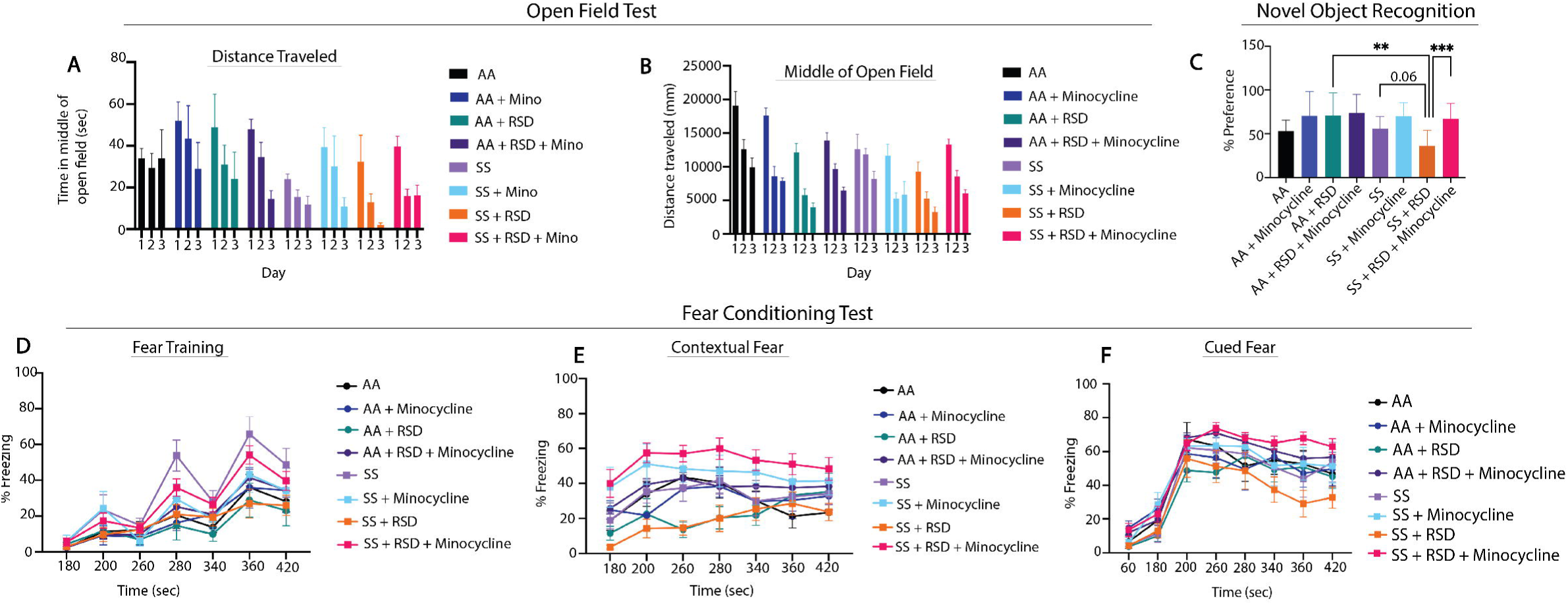
Sickle mice display significant cognitive and neurobehavioral deficits under stress compared to control mice. A)-B) illustrate the results for the open field test, depicting distance traveled through the open field arena and time spent in the middle of the arena. Statistical comparisons were conducted using a two-way ANOVA with Tuckey’s multiple comparisons test. C) depicts preference for the novel object in the NOR test. One-way ANOVA with uncorrected Fisher’s LSD multiple comparisons was conducted to compare object preference between groups. D)-F) illustrate results for the fear conditioning studies (a measure of associative memory). D) represents the training phase, where mice acquired a fear response to an 85 dB tone that was paired with a shock through classical conditioning. E) and F) test the strength of the animals’ conditioned fear response by observing freezing behavior (indicative of fear) after being placed the same environment where the shock had been administered during the learning phase or after hearing the 85dB tone that was associated with the shock, respectively. Freezing behavior was compared between groups using Fisher’s LSD for the training phase and a two-way ANOVA with Holm-Sidak’s adjustment for multiple comparisons for the contextual and cued fear assessments. *p<0.05, **p<0.01, ***p<0.001. Data are presented as mean ± SD.

Furthermore, evaluation of hippocampus-dependent non-associative as well as associative learning and memory, was carried out using NOR and fear conditioning, respectively. In the NOR test Fig. 1C), SS and AA mice had similar percent preference (56.1±13.6% vs. 53.1±12.5%), indicating comparable non-associative memory. However, SS+RSD mice showed evidence of cognitive impairment as demonstrated by lower percent preference (36.1±17.8% SS+RSD vs. 56.1±13.6% SS, *p* = 0.06) compared to SS mice, indicating impaired non-associative memory function. Additionally, we also noted that SS+RSD+minocycline mice had significantly higher percent preference (67.1±17.5% vs. 36.1±17.8%, *p* = 0.0007) compared to SS+RSD mice, suggesting that minocycline treatment led to a sparing of non-associative memory in the treated mice despite exposure to RSD. On the other hand, in the AA group, neither stress nor minocycline treatment were associated with significant changes in cognitive function.

Likewise, the fear conditioning tests (Fig. 1D**-F**), showed that overall, sickle and non-sickle mice irrespective of treatment or exposure disposition, trained similarly during the acquisition phase (Fig. 1D). As shown in Fig. 1E, RSD exposure resulted in significant impairment in contextual (associative) fear memory (evidenced by significantly lower percent freezing) in SS+RSD mice, compared to SS mice. As in Fig. 1C, Fig. 1E, shows that SS+RSD+minocycline mice had significantly better contextual fear memory, compared to SS+RSD mice, p = 0.025 to p < 0.0001 across all time points except at 240s. The abnormal contextual fear memory indicates possible molecular disturbance and/or “overt or covert” lesion of the amygdala resulting from exposure to RSD and its abrogation by minocycline treatment.

Similarly, on cued fear testing (Fig. 1F), there was no significant difference in response from the unperturbed AA or SS mice. However, SS+RSD mice showed significant impairment in hippocampus-mediated cued (associative) fear memory compared to SS+RSD+minocycline mice (p = 0.016 to <0.0001 across different time points. In contrast, neither RSD nor minocycline had significant effects on cognitive function among the AA genotypes.

Next, we evaluated the density of peripheral immune cell infiltrates (known from here on as CD45^+^ bone-marrow derived microglia [BMDM]), Iba1^+^ activated microglia (activation state determined based on morphological features), CD3^+^ T cells, and B220^+^ B cells, in the hippocampus/dentate gyrus (DG). We focused on the hippocampus/DG because of its critical role in cognitive function as well as adult neurogenesis.^31^ In Fig. 2A, we show that overall, SS mice had a higher density of activated microglia (233.6±44.6 cells/mm^2^ vs. 179.3±47.0 cells/mm^2^, *p* ≤ 0.0001) compared to AA mice. Furthermore, SS+minocycline mice had significantly lower density of activated microglia (164.4±56.6 cells/mm^2^ vs. 233.6±44.6 cells/mm^2^, *p* ≤ 0.0001) compared to SS mice. Similarly, SS+RSD mice had significantly higher density of activated microglia (260.2±44.3 cells/mm^2^ vs 233±44.6 cells/mm^2^, *p* ≤ 0.0001) compared to SS mice. Finally, we observed that SS+RSD+minocycline mice had significantly lower density of activated microglia (168.4±45.1 cells/mm^2^ vs. 260.2±44.3 cells/mm^2^, *p* ≤ 0.0001) compared to SS+RSD mice. As shown in Fig. 2A, the results for the comparison within the AA groups, were similar to that described for the SS mice. Additionally, except for the control (non-perturbed group), there were no significant differences between the AA and SS mice based on RSD exposure or minocycline treatment.

**Figure 2.**
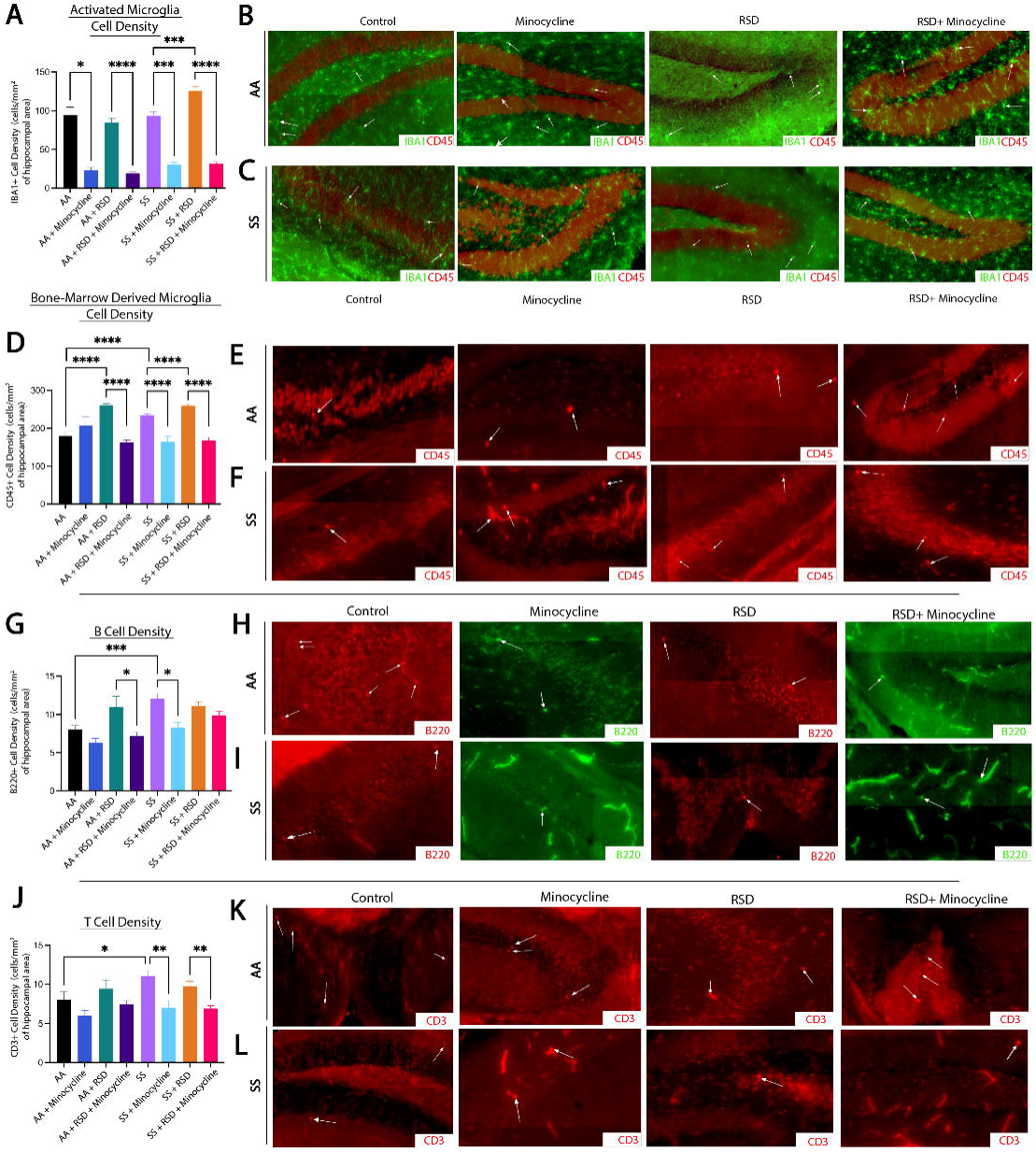
Sickle cell mice exposed to RSD have higher density of activated microglia and B, and T cell infiltrate in the hippocampus while minocycline reduces the density of immune cell infiltrate. A) IBA-1+ activated microglia cell density. B) Representative immunohistochemistry images of IBA-1+ activated microglia in AA control mice, AA mice treated with minocycline, AA mice exposed to RSD, and AA mice exposed to RSD and treated with minocycline. C) Representative immunohistochemistry images of IBA-1+ activated microglia in SS control mice, SS mice treated with minocycline, SS mice exposed to RSD, and SS mice exposed to RSD and treated with minocycline. D) CD45+ bone-marrow derived microglia cell density. E) Representative immunohistochemistry images of CD45+ bone-marrow microglia in AA control mice, AA mice treated with minocycline, AA mice exposed to RSD, and AA mice exposed to RSD and treated with minocycline. F) Representative immunohistochemistry images of CD45+ bone-marrow microglia in SS control mice, SS mice treated with minocycline, SS mice exposed to RSD, and SS mice exposed to RSD and treated with minocycline. G) B220+ B cell density. H) Representative immunohistochemistry images of B220+ B cells in AA control mice, AA mice treated with minocycline, AA mice exposed to RSD, and AA mice exposed to RSD and treated with minocycline. I) Representative immunohistochemistry images of B220+ B cells in SS control mice, SS mice treated with minocycline, SS mice exposed to RSD, and SS mice exposed to RSD and treated with minocycline. J) CD3+ T cell density. K) Representative immunohistochemistry images of CD3+ T cells in AA control mice, AA mice treated with minocycline, AA mice exposed to RSD, and AA mice exposed to RSD and treated with minocycline. L) Representative immunohistochemistry images of CD3+ T cells in SS control mice, SS mice treated with minocycline, SS mice exposed to RSD, and SS mice exposed to RSD and treated with minocycline. Cell density was compared between groups with a one-way ANOVA and Fisher’s LSD multiple comparisons test. *\*p<0.05, **p<0.01, ***p<0.001. Data are presented as mean ± SD*.

Results of the examination of the contribution of peripheral immune cell (CD45^+^) infiltrate to the observed cognitive deficit and impact of minocycline treatment is shown in Fig. 2D. There was no significant difference between SS and AA mice with respect to the density of CD45^+^ BMDM cells (93.7±45.8 cells/mm^2^ vs. 94.8±50.6 cells/mm^2^) in the hippocampus/DG. However, both AA+RSD and SS+RSD mice had significantly higher density of CD45^+^ BMDM compared to their non-perturbed controls, with SS+RSD mice having a significantly higher density of CD45 ^+^ BMDM compared to AA+RSD mice (125.4±70.3 cells/mm2 vs. 84.8±60.5 cells/mm2, *p* < 0.0001).

Additionally, AA+minocycline mice had a significantly lower CD45^+^ BMDM compared to AA mice (23.4±10.0 cells/mm^2^ vs. 94.8±50.6 cells/mm^2^, *p* = 0.013). Similarly, AA+RSD+minocycline mice had a significantly lower density of CD45^+^ BMDM (19.1 ± 10.4 cells/mm^2^ vs. 84.8±60.5 cells/mm^2^, *p* < 0.0001) compared to AA+RSD mice. Following in a similar pattern, SS+minocycline mice had significantly lower density of CD45^+^ BMDM (30.0 ± 12.5 cells/mm^2^ vs. 93.7±45.8 cells/mm^2^ *p* = 0.0002) compared with SS mice, and SS+RSD mice having a significantly higher density of CD45^+^ BMDM (125.4±70.3 cells/mm^2^ vs. 93.7±45.8 cells/mm^2^, *p* = 0.0004) compared to SS. We also observed that SS+RSD mice had significantly higher density of CD45^+^ BMDM (125.4±70.3 cells/mm^2^ vs. 84.8±60.5 cells/mm^2^, *p* < 0.0001) compared to AA+RSD mice. On the other hand, SS+RSD+minocycline mice had significantly lower density of CD45^+^ BMDM (31.8±14.9 cells/mm^2^ vs. 125.4±70.3 cells/mm^2^, *p* < 0.0001) compared to SS+RSD mice.

Furthermore, we quantified the density of B cells (B220^+^) and T cells (CD3^+^) and this is shown in Fig. 2G. Of note, AA+RSD mice had significantly higher density of B cells (11.0±3.7 cells/mm^2^ vs. 7.2±3.0 cells/mm^2^, *p* = 0.037) compared to AA+RSD+minocycline mice. There was also a slightly lower density of B cells in AA+minocycline mice (6.3±1.9 cells/mm^2^ vs. 8.1±1.9 cells/mm^2^) compared to AA mice, however, this was not statistically significant. Suggesting that minocycline may be suppressing B cell mediated neuroinflammation by limiting peripheral immune cell infiltration into the brain. Similarly, SS mice had significantly higher density of B cells (12.1±3.4 cells/mm^2^ vs. 8.1±1.9 cells/mm^2^, *p* = 0.0009) compared with AA mice and SS+minocycline mice (12.1±3.4 cells/mm^2^ vs. 8.3±2.3 cells/mm^2^ *p* = 0.012). Surprisingly, exposure of SS mice to RSD with or without minocycline treatment did not result in a significant change in B cell density, contrary to our observation among the AA mice. This result suggests that B cell infiltration might play a smaller role in RSD-induced neuroinflammation as an underlying mechanism for the development of cognitive deficits in SCD.

Further analysis as shown in Fig. 2J, indicates that SS mice had significantly higher T cell density (11.1±3.8 cells/mm^2^ vs. 8.1±3.8 cells/mm^2^, *p* = 0.032) compared to AA mice and AA+minocycline mice had lower density of T cells compared to AA mice (6.0 ± 2.2 cells/mm^2^ vs. 8.1 ± 3.8 cells/mm^2^, *p* = 0.032). There was not statistically significant difference in the comparison of T cell density in AA+RSD mice to that of AA+RSD+minocycline mice (7.5±2.9 cells/mm^2^ vs. 9.4±2.9 cells/mm^2^). Among sickle cell mice, we observed that SS+minocycline mice had a significantly lower T cell density (7.0±2.8 cells/mm^2^ vs. 11.1±3.8 cells/mm^2^, *p* = 0.005) compared to SS mice, and no significant difference in T cell density between SS mice and SS+RSD mice (11.1±3.8 cells/mm^2^ vs. 9.8±3.4 cells/mm^2^). However, SS+RSD+minocycline mice had a significantly lower T cell density (6.9±2.2 cells/mm^2^ vs. 9.8±3.4 cells/mm^2^, *p =* 0.0018) compared to SS+RSD mice. This indicates a possible but slightly lesser role for T cells in cognitive impairment in SCD in the setting of exposure to social stress.

Given the reported role of neurogenesis in social stress-induced neuroinflammation and cognitive deficits,^18, 32^ we quantified and compared the densities of neural progenitor cells (NPCs; DCX^+^), adult-born neurons (DCX^+^NeuN^+^) and “newly formed” astrocytes (DCX^+^GFAP^+^) in the dentate gyrus and report our findings in Fig 3. As shown in Fig. 3A, we observed a higher density of NPCs in AA+minocycline mice (20.8±4.8 cells/mm^2^ vs. 17.1±5.7 cells/mm^2^) compared to AA mice, while the NPC density was lower among AA+RSD mice (13.3±3.6 cells/mm^2^ vs. 17.1±5.7 cells/mm^2^ p = 0.06) compared to AA mice. Additionally, AA+RSD mice had significantly lower NPC density compared to AA+RSD+minocycline mice (13.3±3.6 cells/mm^2^ vs. 18.7±5.3 cells/mm^2^ p < 0.0001). We also observed that SS mice had slightly lower NPC density (15.3±5.3 cells/mm^2^ vs. 17.1±5.7 cells/mm^2^) compared to AA mice and, SS+minocycline mice had a significantly lower NPC density (14.1±4.4 cells/mm^2^ vs. 20.8±4.8 cells/mm^2^, *p* = 0.006) compared to AA+minocycline mice. Furthermore, as seen in the AA groups, SS+RSD mice had a significantly lower NPC density (11.9±3.9 cells/mm^2^ vs.15.8±5.6 cells/mm^2^ (*p* = 0.005) compared to SS+RSD+minocycline mice. We also noted that SS+RSD+minocycline mice have essentially the same NPC density as SS mice, indicating that minocycline might be limiting the gliogenic shift that seems to result from exposure to RSD (see Fig. 3B and **Supplementary** Fig. 1), leading to development of cognitive deficit.

**Figure 3.**
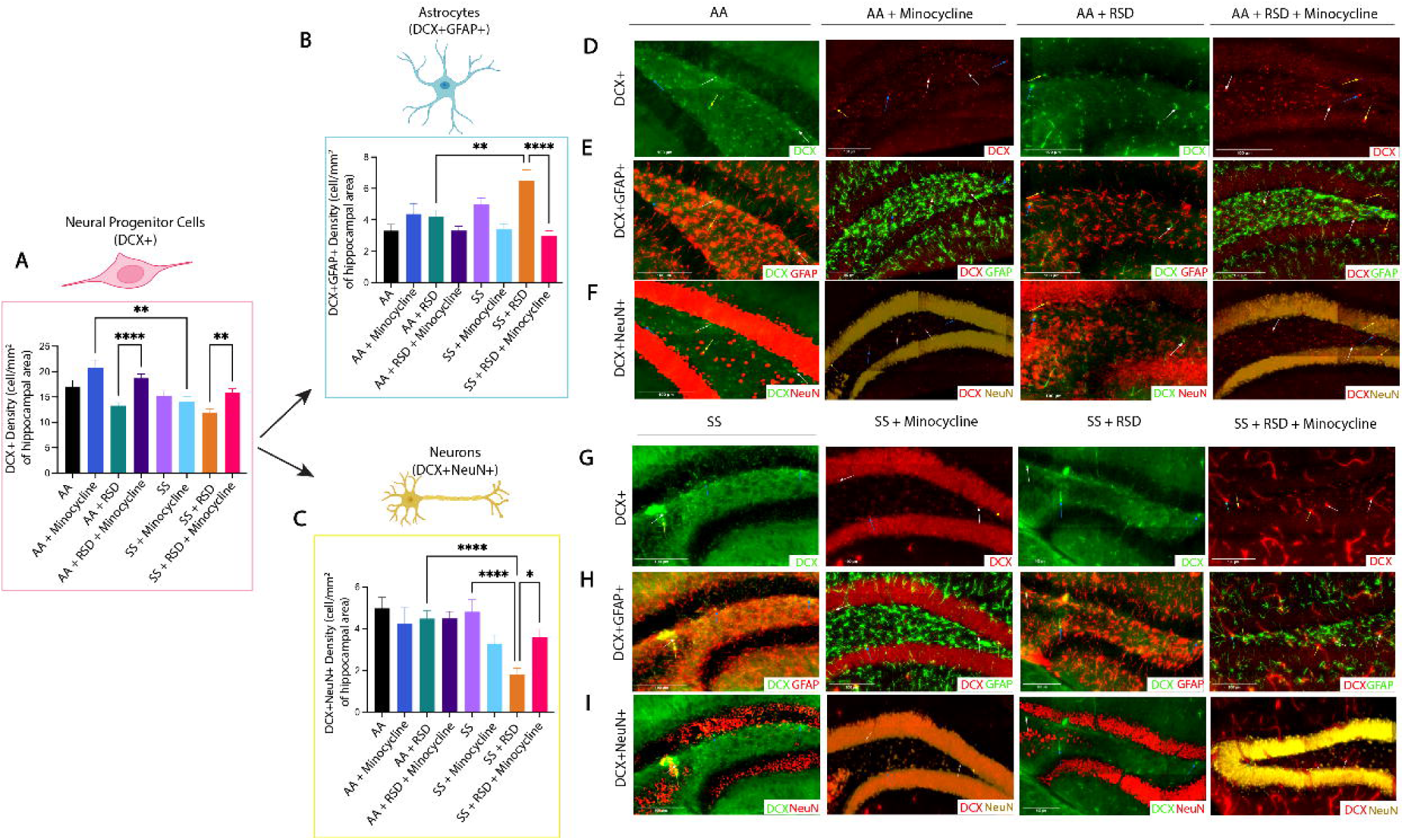
Sickle cell mice exposed to RSD have higher density of DCX^+^GFAP^+^ astrocytes while having decreased densities of DCX^+^ neural progenitor cells and DCX^+^NeuN^+^ neurons in the hippocampus. A) DCX^+^ neural progenitor cell density. B) DCX^+^GFAP^+^ astrocyte cell density. C) DCX^+^NeuN^+^ neuron cell density D) Immunohistochemistry of DCX^+^ neural progenitor cells in AA mice. E) Immunohistochemistry of DCX^+^GFAP^+^ astrocytes in SS mice. F) Immunohistochemistry of DCX^+^NeuN^+^ neurons cells in AA mice. G) Immunohistochemistry of DCX^+^ neural progenitor cells in SS mice. H) Immunohistochemistry of DCX+GFAP+ astrocytes in SS mice. I) Immunohistochemistry of DCX^+^NeuN^+^ neurons in SS mice. Statistical comparisons were performed with a one-way ANOVA with Fisher’s LSD multiple comparisons test. *\*p<0.05, **p<0.01, ****p<0.0001. Data are presented as mean ± SD. ****p<0.0001*.

Furthermore, as shown in Fig. 3B, SS mice had a higher density of newly formed astrocytes (5.0±1.9 cells/mm^2^ vs. 3.3±1.9 cells/mm^2^) compared to AA mice, while SS+minocycline mice had a lower density of new astrocytes (3.4±1.5 cells/mm^2^ vs. 5.0±1.9 cells/mm^2^) compared to SS mice. On the other hand, SS+RSD mice had a significantly higher density of new astrocytes (6.5±3.9 cells/mm^2^ vs. 4.2±2.7 cells/mm^2^, *p* = 0.0011) compared to their matched AA+RSD counterparts, as well as when compared to SS+RSD+minocycline mice (i.e., 6.5±3.9 cells/mm^2^ vs. 2.9±2.3 cells/mm^2^, *p* < 0.0001). We also noted that AA+RSD mice had a higher density of new astrocytes (4.2±2.7 cells/mm^2^ vs. 3.3±1.8 cells/mm^2^) compared to AA mice, but this was not statistically significant. Taken together, this indicates that the exposure to RSD alone might be shifting the differentiation of NPCs towards astrocytes. And that treatment with minocycline reduces that shift as seen in the SS+RSD+minocycline and AA+RSD+minocycline mice when compared to their untreated but stressed counterparts.

Results of the quantification of the density of adult-born neurons (DCX^+^NeuN^+^) is shown in Fig. 3C and **Supplementary** Fig. 1. Overall, we show that AA mice irrespective of treatment or RSD status, had similar adult born neuron densities. Among the SS mice, SS+RSD mice had significantly lower adult-born neuron density compared with SS mice (1.8±1.7 cells/mm^2^ vs. 4.8±2.9 cells/mm^2^, p < 0.0001), as well as compared to AA+RSD mice (1.8±1.7 cells/mm^2^ vs. 4.5±2.4 cells/mm^2^, p < 0.0001). SS+RSD mice also had a significantly lower density of adult-born neurons compared to SS+RSD+minocycline mice (1.8±1.7 cells/mm^2^ vs. 3.6±2.6 cells/mm^2^ p = 0.0031). **Supplemental** Fig. 1 provides the percentage distribution of these cells (DCX^+^GFAP^+^ and DCX^+^NeuN^+^) as a percentage of the total NPCs (DCX^+^) counted.

The result of bulk RNA sequencing and analysis (**Supplementary** Fig. 2A and 2B) including gene-set enrichment analysis is presented in the results section of the online supplementary material. Also included in the online supplementary material are the results of lipidomics analysis (**Supplementary** Fig. 3A and 3D).

## Discussion

In this study, we sought to understand how RSD affects cognitive function (with or without treatment) in humanized Townes sickle mice, compared to controls (treated and untreated) mice. Our findings presented above and the online supplemental materials, supports our stated hypothesis, and shows that a social stressors (RSD), impairs cognitive functions in sickle mice, similar to what was described among children with SCD, by King et al.^8, 9^ It also provide some mechanistic insight, in showing that neuroinflammation and possibly depression of neurogenesis (Fig. 3), with a shift towards astrogliogenesis (**Supplementary** Fig. 2) may be among the underlying mechanisms. In our prior work, we showed that 13 months old Townes sickle mice had more severe cognitive and neurobehavioral deficits and abnormal neuroplasticity.^33^ The findings from that study, motivated this work in understanding why children with SCD living in a socially stressful environment have more severe manifestation of cognitive deficit. Thus, we additionally showed that minocycline treatment alleviates neuroinflammation and neurogenesis and leads to better cognitive and neurobehavioral functions and well as improvement in relevant molecular and cellular phenotypes.

It is known that individuals with SCD experience cognitive and neurobehavioral (anxiety and depression) deficits observed in early childhood, adolescents and adults.^34, 35^ We saw sickle mice exhibit cognitive and neurobehavioral deficits after being exposed to social stress, recapitulating what was described in children with SCD. In these children, it was shown that the presence of cognitive deficit was associated with “biological factors” such as severity of anemia and presence of silent cerebral infarct (SCI) or stroke.^36–38^ However, King et al^8, 9^ demonstrated more severe evidence of cognitive deficit in children without SCI, but who were exposed to social stress in the forms of low parental socioeconomic status. This and the report by Andreotti et al.^7^, were essentially recapitulated in our study, which showed one or more mechanisms underlying the development of cognitive deficit in children with SCD.

In our study, evaluating the hippocampus and dentate gyrus revealed the presence of evidence of neuroinflammation in sickle mice at baseline, i.e., without exposure to RSD. We noted that sickle mice exposed to stress had higher densities of “activated microglia” and CD45^+^ bone-marrow derived microglia compared to control mice or sickle cell sickle cell mice exposed to stress and treated with minocycline. These findings are of particular interest as increased microglia activation or overactive microglia undergoes phenotypical and functional changes often resulting in increased pro-inflammatory cytokine secretion and increased phagocytosis. These activities have been shown to be involved in the mechanism of cognitive impairment and causes neurobehavioral changes (anxiety and anhedonia).^39–41^ Additionally, studies have shown that peripheral mononuclear cell or bone-marrow derived microglia, infiltrate the brain parenchyma after psychological stress, and further leads to neuroinflammation, anxiety and memory deficit.^42^ Taken together, this suggests that social stress promotes peripheral immune cell infiltration regardless of genotype, but more so in sickle cell which is already a pro-inflammatory state. In our study, we did not see increased lymphocyte densities in sickle mice exposed to stress, however we did note that sickle mice overall, without exposure to RSD, had higher T and B cell densities as well as a higher density of bone-marrow derived microglia compared to AA control mice, supporting our assertion of a background neuroinflammation in SCD, which was accentuated by exposure to RSD, leading to cognitive deficit. Additionally, recent studies have reported that B cells contribute to neuroinflammation via peripheral immune mechanisms, through the production of pro-inflammatory cytokines and antibodies while effector T cells interaction with microglia can further promote inflammation.^43, 44^ This may explain why we observed a higher density of T cells with RSD exposure, but did not observe a higher density of B cells. Furthermore, the presence of a higher density of peripheral immune cells in the brain, in sickle cell mice indicates their possible role in SCD-related neuroinflammation even in the absence of social stress. We did not adequately examine the presence of T or B cells for instance, in our prior study, however it is conceivable to assume they were involved in our observation.^33^ Overall, these findings illustrate the potential cellular mechanisms that contribute to cognitive deficits in sickle cell mice exposed to stress and could underly the observation among SCD patients exposed to social stress such as lower individual or parental socioeconomic status.

Chronic social stress modulates neurogenesis by decreasing neuron proliferation, resulting in modifications to hippocampal synaptic signaling and plasticity.^45^ However, the effect of stress on neurogenesis in SCD is still unknown. In our study, we noted that neural progenitor cells (NPCs) in the dentate gyrus of sickle mice exposed to RSD, shifted more (in their differentiation) towards astrogliogenesis, as opposed to mature neurons. It has been documented that minocycline improves neurogenesis and mitigates the gliogenic effect of inflammatory cytokines on NPCs. ^46–49^ This was also observed in our study, where we noted that SS+RSD+minocycline mice had significantly, higher density of adult born neurons, lower density of new astrocytes and lower density of proinflammatory cells in the hippocampus/DG compared to SS+RSD mice. Intriguingly, minocycline treatment of AA mice led to increased NPC density as well, but not on density of adult born neurons. The analysis of bulk RNA sequencing and lipidomics supported our other findings and is presented as well as discussed in the online supplementary material.

In conclusion, we have attempted to show some of the underlying mechanisms of how RSD affects cognitive deficits, akin to those described in children with SCD exposed to social stress. We showed that the development of cognitive deficit is in part driven by “activation” of resident immune cells and/or infiltration of peripheral immune cells, astrogliogenesis, changes to lipid metabolism and the transcriptome. Finally, we demonstrated that treatment with minocycline (which is anti-neuroinflammatory and a sphingomyelinase inhibitor), mitigated the presence of cognitive deficit possibly by blocking neuroinflammation, shifting NPCs towards neurogenesis and as well as supporting a favorable lipidomics and transcriptomic profile that supports neurogenesis and synaptogenesis as well as synaptic plasticity, but is anti-excitotoxic, and anti-neuroinflammatory.

## Acknowledgement

This was supported in part by the following awards: S’D.A.D was supported by the National Institutes of Health (T32 GM145735). O.T.G. was supported by the American Society of Hematology Scholar Award and the Parker B. Francis Fellowship in Pulmonary Research Award.

H.I.H. is supported by grants from the National Institutes of Health (R01 HL138423, R01 HL156024 and 3R01 HL138423-05S).

## Author Contribution

S’D.A.D, O.T.G and S.V wrote the initial draft of the manuscript, H.I.H designed the study, carried out some of the experiments, R.Z.H and A.S.S performed immunohistochemistry and carried out microscopy imaging and K.P, R.Z.H and H.I.H performed analyses of immunohistochemistry images. L.N performed alignment and analysis of RNA sequencing data, M.S and A.G.J, S.V and H.I.H performed bioinformatics and gene-set enrichment analysis. S’D.A.D performed lipidomics assay and E.A.B, S’D.A.D, H.I.H and V.S performed lipidomics analyses. H.I.H and O.T.G performed critical review of the manuscript. All authors approved the manuscript for submission.

