## Supplementary Text for "Neuroinflammation underlies the development of social stress induced cognitive deficit in sickle cell disease"

**Methods**

*Repeated Social Defeat (RSD)*: RSD was performed as previously described in the literature [1] to model sociological stress. In brief, an aggressive male intruder mouse was introduced into cages of established male cohorts for two hours, between 5:00 P.M. and 7:00 P.M. for six consecutive nights. Each cohort contained three HbSS or HbAA mice. If the intruder did not initiate an attack, or was attacked by a resident mouse, within 5–10 minutes, a new intruder would be introduced. During each 2-hour cycle, we observed submissive behavior, including upright posture, fleeing, and crouching, to ensure defeat of the resident mice. At the end of the 2-hour period, the intruder mouse was removed from the cage and the residents were undisturbed until the beginning of the next 2-hour cycle on the following day. No intruder mice were introduced to control mice cages.

*Minocycline treatment*: Minocycline (Sigma-Aldrich) was administered orally in drinking water, and its effects on neuroinflammation, neuronal morphology and plasticity, and brain lipidomic on genotype were examined. Minocycline solutions were prepared fresh daily and administered in opaque sipper bottles because minocycline is photosensitive. To adjust administration dosages to 90 mg/kg based on drinking pattern, mice and water bottles were weighed daily. Oral minocycline treatment began 1 day before the beginning of RSD and was terminated on the last day of RSD, which is approximately 24 hours before cognitive and neurobehavioral evaluation.

*Open field test and novel object recognition:* The open field test and novel object recognition (NOR) test were conducted as part of the same experiment. The open field test measures anxiety-like and depressive behaviors in mice,[2] while the NOR test is used to evaluate hippocampus-based episodic memory and learning.[3-5] Both tests were conducted in an open field arena made **of** plexiglass measuring 12” x 12” x 15” (L x W x H). The details of how these behavioral tests are conducted have been published by our lab here[6] as well as by others.[3, 5, 7-9] Briefly, mice were allowed 2-3 days of habituation, where they were allowed to explore the arena for 5 minutes every day. On the 3rd or 4th day, the mice were trained by placing two identical test objects in the enclosure and allowing the mice to explore them for 5 min. Following training, the mice were returned to their home cages for about 30 min. After this delay, the learning and memory of the mice were tested by replacing one of the test objects used during training with a novel object in the open field.

*Fear conditioning tests:* The fear conditioning tests are used to evaluate associative memory in mice by measuring their ability to form and retain associations between aversive experiences and environmental cues. The tests were performed over a 3-day period, as previously described.[4] Briefly, on the first day, mice were placed in the fear conditioning chamber and allowed to explore for 3 minutes. Afterwards, mice were presented with 3 pairs of conditioned and unconditioned stimuli. In each pair, the conditioned stimulus (CS) consisted of a 20-second 85 dB tone, and the unconditioned stimulus (US) consisted of a 2-second 0.5 mA electric shock to the footpad. One minute after the last CS-US pair was presented, the mice were returned to their home cages. On the second day, mice underwent a contextual fear conditioning test. The animals were returned to the fear conditioning chamber, but this time no shock was delivered. Freezing behavior was assessed over a 9-minute period. The percent of each minute that mice spent frozen was recorded. On the third day, mice underwent a cued fear conditioning test. The animals were placed in a new environment and allowed to explore for 2 minutes. Afterwards, the CS was presented every minute for 9 minutes, freezing behavior was recorded every minute. In both tests, freezing indicates a memory for either the context in which the shock was delivered or the association between the tone and the shock. More freezing indicates better associative learning and memory.[4]

*Immunohistochemistry and digital image analysis for IBA-1 and CD45:* After the cognitive and behavioral testing, mice were brough back to their home cages and sacrificed the next day. Brain samples were collected after transcardial perfusion with sterile phosphate buffered saline (PBS), pH 7.4, followed by 4% formaldehyde in PBS. Brains were post-fixed in 4% formaldehyde for 24 h and then transferred to 0.1% sodium azide in PBS if not immediately sectioned and stored at 4°C. Fixed brains were sectioned (50µm) using a vibratome (1200S Leica Microsystems). Brain regions within the hippocampus were identified by reference markers in accordance with the stereotaxic mouse brain atlas.[7] To label for IBA-1 or CD45, sections that were stored in azide were washed in PBS and incubated overnight at room temperature, in primary antibody cocktail containing rabbit anti-mouse Iba-1 (1:1000; Wako Chemicals), rat anti-mouse CD45 (1:500; Abcam) and guinea pig anti-mouse NeuN, diluted in antibody diluting solution (containing 0.1% azide, 2% Triton-x-100 and 10% normal goat serum in PBS). Then sections were washed in PBS and incubated with a fluorochrome-conjugated secondary antibody (Alexa Fluor 488 and 594 and 750 respectively). Sections were then mounted on glass slides, cover-slipped with Fluoromount G (Beckman Coulter) and stored at 4°C after drying. Images (z-stacks) of the dentate gyrus on either side, in all 3 fluorochrome color channels, were acquired at 20X magnification using an SP8 confocal microscope (Leica Microsystems), projected using maximum intensity projection and then analyzed using NIH ImageJ software. CD45, cells with positive labeling will be counted in each dentate gyrus section, the dentate gyrus is a component of the hippocampus. IBA-1 labeling were analyzed using established and published digital image analysis system.[8] In brief, a threshold for positive labeling was determined for each image that included all cell bodies and processes but excluded background staining. Data were processed by densitometric scanning of the threshold targets using NIH ImageJ software. The proportional area were reported as the average percentage area in the positive threshold for all representative images.[10]

*Sphingolipid Extraction and LC-MS/MS Analysis:*

The details of the protocol used for sphingolipid extraction has been described elsewhere.[11] Briefly, hippocampal and cortical tissues were homogenized in PBS. Each homogenate was divided into two aliquots. The first aliquot underwent sphingolipid extraction, while the second aliquot was used for total protein quantification with the BCA protein assay. The first aliquot was further divided between long-chain bases (LCBs) and complex sphingolipids (CSLs) extractions. Samples designated for LCB extraction were suspended in 2:1 methanol:methylene chloride solution, while the samples designated for CSL extraction were suspended in 2:1 methanol:chloroform. Internal standard mixture (Avanti Polar Lipids) was then added to each sample, and samples were incubated overnight. Afterwards, the CSL samples underwent base catalysis for 2 hours before being neutralized. The LCBs samples were transferred to new glass tubes, leaving behind tissue debris, and 2:1 methanol:methylene chloride extraction solvent was added to the original glass tubes. Then, tubes were centrifuged to collect residual LCBs from the debris, and the extraction solvent (containing lipids) was transferred to the new glass tube. For the CSLs samples, the extraction solvent consisted of water added to the 2:1 methanol:chloroform mixture, allowing for aqueous-organic phase separation. The CSL samples were centrifuged, and the bottom organic layer (containing lipids) was transferred to new glass tubes, leaving behind tissue debris. The extraction solvent was again added to the residual tissue debris, and the samples were centrifuged a second time. The bottom organic phase was once again collected. Finally, the organic solvents from both LCB and CSL samples were subsequently removed by vacuum drying overnight in a Savant SpeedVac (Thermo Fisher). The dried lipids were stored in a -20 C freezer until analysis.

*Analysis of RNA Sequencing, Data Analysis and GO Analysis:*

Directional polyA RNA-seq was performed by the Genomics, Epigenomics and Sequencing Core at the University of Cincinnati using established protocols as previously mentioned.[12, 13] To summarize, the quality of total RNA was QC analyzed by Bioanalyzer (Agilent, Santa Clara, CA). To isolate polyA RNA for library preparation, NEBNext Poly(A) mRNA Magnetic Isolation Module (New England BioLabs, Ipswich, MA) was used with 500 ng good quality total RNA as input. The polyA RNA was enriched using SMARTer Apollo automated NGS library prep system (Takara Bio USA, Mountain View, CA). Next, NEBNext Ultra II Directional RNA Library Prep kit (New England BioLabs) was used for library preparation under PCR cycle number of 9. After library QC and quantification via real-time qPCR (NEBNext Library Quant Kit, New England BioLabs), individually indexed libraries were proportionally pooled and sequenced using NextSeq 550 sequencer (Illumina, San Diego, CA) under the sequencing setting of single read 1x85 bp to generate about 25M reads. After sequencing, fastq files for downstream data analysis were automatically generated via Illumina BaseSpace Sequence.

The raw sequencing reads were aligned to GRCm39 (mm39) using STAR.[14] The number of reads that were aligned to each gene was obtained while mapping using the GENCODE gene annotation. Then we used edgeR[15] to identify differentially expressed genes. For each genotype ("AA" or "SS") in each tissue (Hippocampus, cortex), we included all samples in a model and compared the two "exposure" groups (RSD vs. No RSD) and then "treatment" groups (minocycline treatment vs. no minocycline treatment) in a pairwise fashion. We also compared across genotype; however, we ensured that only biologically meaningful comparisons were performed.

*Functional enrichment analysis and visualization*:

We used ToppFun[16] and ToppCluster[17] applications to perform functional enrichment analysis of differentially expressed genes (DEGs) in various groups. Genes differentially expressed in different comparisons were input into the ToppFun/ToppCluster for gene ontology (GO) and biological pathway annotation, with Benjamini and Hochberg’s correction for significance testing set at p value <0.05. To visualize the enriched biological processes and pathways, we used Cytoscape.v3.0.2[18] (for network representation) and Morpheus (https://software.broadinstitute.org/morpheus/; for heatmap representation) application.

The raw and processed RNA sequencing data along with the metadata have been uploaded to GEO with the accession **GSE252778**.

### Results

In **Supplementary Fig. 1**, we assessed the percent distribution of neural cells to evaluate how RSD and minocycline treatment influence neural cell distribution. When looking at the percentage of $DCX^+$ neural progenitor cells (NPCs) baseline AA mice had 50.1% NPCs. AA+minocycline mice had approximately 59.9% NPCs. We observed a lower percent of NPCs in AA+RSD mice (36.5%); however, AA+RSD+minocycline mice had an increase in NPCs (57.7%). SS mice had a lower percentage of NPCs (35.1%) compared to their AA counterparts and treating sickle mice with minocycline increased NPCs percent to 51.2%. SS+RSD mice had the lowest percent among all the groups, with only 29.3% NPCs. Similar to AA mice with the same treatment, treatment of sickle mice exposed to RSD with minocycline resulted in an increase of NPCs (58.7%). Next, we assessed $DCX^+NeuN^+$ neuron distribution. In AA control mice, we saw a 30.0% neuron distribution; however, treatment with minocycline decreased the neuron percent to 19.4%. AA+RSD mice had 33.4% neurons, while treatment with minocycline decreased this to 23.9%.

Comparable to AA control mice, SS mice had 30.0% neurons. This fraction was lower in SS+minocycline mice (23.7%). SS+RSD mice had 17.6% of neurons, and treatment with minocycline increased this to 20.6% of neurons. Lastly, we looked at $DCX^{+}GFAP^{+}$ astrocyte distribution. We observed that baseline AA mice had a similar percentage of astrocytes with AA+minocycline mice. AA+RSD mice had 30.1% astrocytes; however, a decrease of astrocytes (18.4%) was noted when AA+RSD mice were treated with minocycline. SS control mice had 34.9% astrocytes, while SS+minocycline mice had 25.1% astrocytes. SS+RSD mice had 53.1% astrocytes, and treatment with minocycline reduced this to 20.7%. This data suggests that RSD decreases NPCs in both AA and SS mice while increasing astrocyte, Additionally, treating mice exposed to stress using RSD caused a decrease in astrocytes and an increase in NPCs. Minocycline treatment may prevent NPC differentiation into astrocytes.

Using the results of our bulk RNA sequencing analysis, we performed gene set enrichment analysis (GSEA) to identify the pathways or processes that may underly RSD-linked cognitive deficit and neuroinflammation in SCD. Most of the differentially expressed gene sets in the cortex were involved in cognitive function, synaptic structures, neuronal signaling, and inflammation (**Supplementary Fig. 2A**). Differences between sickle cell and AA healthy control mice were evident both at baseline and after RSD exposure. We observed that genes connected with blood-brain barrier dysfunction, depressive disorders, and inflammation (*CCR7*, *FOXF1*) were enriched in SS relative to AA mice. SS+RSD mice show enrichment for genes related to neurodegenerative disease processes (*CTNNB1,*[19] *CSF1R*, *VCP* [20-22]), while sirtuins, which prevent aging and neurocognitive diseases,[23, 24] were less enriched. In contrast, no significant changes in gene expression were observed in AA+RSD group. Additionally, *LDLR* expression (associated with long-term memory) was downregulated in the SS+RSD group compared to the AA+RSD group. Together, these findings support our hypothesis that SS mice might have greater susceptibility to the effects of social stressors such as RSD.

Furthermore, we observed that SS+RSD+minocycline mice showed enrichment for genes related to processes governing synaptic structure and plasticity (*BDNF*, *ENTPD1*), while genes associated with overall cell signaling, immune infiltration and lipid membrane trafficking (*Adora1, ABDH6*, *Akt1/2*) were downregulated. Notably, excitatory signaling through serotonin receptors (5-HTR 4, 6, and 7), glutamatergic, and dopaminergic synapses was decreased, highlighting a possible role for minocycline in preventing stress-linked excitotoxicity and neurodegeneration.[25, 26] Taken together, our results suggest that minocycline may help prevent imbalances in synaptic activity and functional decline in sickle mice exposed to stress. Finally, pathways linked to inflammation, gliogenesis, and neuronal death were also less enriched in SS+RSD+minocycline animals, while sirtuin related pathways and processes were enriched.

In the hippocampus, similar trends were observed (**Supplementary Fig. 2B**), as we saw that genes related to abnormal cerebral vasculature, blood-brain barrier dysfunction, and inflammatory processes were more enriched in SS compared to AA mice. Furthermore, in SS+RSD mice, genes related to brain development (*MAOB*) and neurodegeneration were significantly enriched compared to AA+RSD mice. Additionally, compared to AA+RSD animals, SS+RSD mice showed significant enrichment of genes negatively associated with forebrain morphogenesis, and neurogenesis, and positively associated with inflammation. Together, these findings suggests that exposure to RSD/social stressors have a negative impact on the brain, including structural remodeling, and this appears to be more pronounced in the sickle cell mice. As was observed in the cortex, minocycline treatment was associated with a unique enrichment signature in sickle mice exposed to stress. For instance, genes that are negatively associated with remodeling of cerebral structures and blood-brain barrier integrity, brain development and neurogenesis, and positive associated with inflammation were all down-regulated in SS+RSD+minocycline mice.

Together, these results support minocycline's function in preventing neuroinflammation, evidence of neurodegeneration, and cognitive deficit in sickle cell mice exposed to stress and suggests that these might underly the mechanism of social stress related cognitive deficit in SCD.

Because sphingolipids play important roles in neurological function and immune signaling, we sought to understand whether they may be connected with neuroinflammation, and social stress induced to cognitive deficit in SCD. GSEA analysis was performed to evaluate enrichment of sphingolipid-related pathways in the cortex (**Supplementary Fig. 3A**) and hippocampus (**Supplementary Fig. 3C**), while we used liquid chromatography-mass spectrometry (LC-MS) to quantify the concentrations of sphingolipids found in these two brain regions (**Supplementary Figs. 3B, 3D**).

In the cortex (**Supplementary Fig. 3A**), AA and SS mice differ significantly in their gene expression profiles with and without exposure to stress. The top processes enriched in SS mice compared to their AA counterparts include cellular responses to lipid stimuli and regulation of lipid kinase activity, while processes downregulated in SS mice include regulation of lipid synthesis, metabolism, transport, and storage. In SS+RSD+minocycline mice, we observed substantial changes in enrichment patterns for sphingolipid-related pathways and processes. For instance, biological processes governing metabolism, transport, and storage of various lipid species, including sphingolipids, were downregulated in the SS+RSD+minocycline mice compared to the SS+RSD mice. Overall, genes associated with sphingolipid signaling and metabolism pathways were also downregulated. In particular, SS+RSD+minocycline mice showed lower expression of genes associated with ceramide (Cer) metabolism, including de novo cer synthesis (*Elovl4*, *Slc1a4*),[27] degradation of sphingomyelin (SM) into cer via the salvage pathway (*SMPD1*), and synthesis of ceramide-derived sphingolipids, including sphingosine-1-photphate (S1P) (via *Sphk1*) and complex gangliosides (via *ST3GAL2*).[28]

We also examined the results of the LC-MS analysis to identify a link between the pathway enrichment/gene expression and lipidomics profile and levels in the cerebral cortex or hippocampus. In **Supplementary Fig. 3B**, we showed that AA mice had higher levels of Cer, and SM and lower levels of hexosylceramide (HexCer), Sphingosine (Sph) and S1P compared to AA+RSD mice. Interestingly, SS mice with or without exposure to RSD, had the highest level of Cer of the two genotypes, compared to their AA counterparts. Also, SS mice had higher levels of HexCer, SM, Sph, and S1P compared to SS+RSD mice. This later point may indicate that exposure to RSD/social stress alters the sphingolipid profile by potentially decreasing enzymatic activity in the sphingolipid and thus sphingomyelin biosynthetic and degradation pathways. Minocycline treatment had a significant impact on sphingolipid profile. As stated earlier, SS+RSD mice had low levels of SM, Sph, and S1P, while AA+RSD mice had high levels of Sph and S1P. However, we saw that AA+RSD+minocycline mice had increased levels of Cer, HexCer, SM, and LSM and low levels of Sph and S1P compared to AA+RSD mice. Similarly, SS+RSD+minocycline mice had higher levels of SM, Sph and S1P and a lower level of LSM compared to SS+RSD. Overall, these findings suggest that minocycline could restore sphingolipid enzymatic activity perturbed by social stress, and thus could be one mechanism of its benefit.

Likewise, GSEA and LC-MS analyses of the hippocampal tissue showed that genes involved in a few but critical biological processes were differentially expressed between AA and SS mice, as well as between SS+RSD and SS mice (**Supplementary Fig. 3C**). Specifically, we observed that compared with SS+RSD mice, SS+RSD+minocycline mice showed downregulation of processes related to metabolism, synthesis, and transport of lipid species (including sphingolipids), as well as responses to lipid stimuli. Notably, several genes responsible for inhibiting (1) de novo ceramide synthesis (*ORMDL2*/*ORMDL3*),[29] (2) breaking down lysosomal sphingomyelin to ceramide (*SMPD1*), (3) synthesizing gangliosides from ceramide (*ST3GAL2*/*ST3GAL3*),[28] and (4) converting S1P to sphingosine (*PLPP3*), were significantly less enriched compared to SS+RSD.[30] We noted that some of the genes included in the response to lipid stimuli process (*CD38, CX3CR1*, and *TLR2*) and thus down regulated in the SS+RSD+minocycline mice, also encode some of the surface receptors found on microglia and lymphoid cells;[31-34] hence, lower expression of these genes in the treated mice may explain the reduced evidence of neuroinflammation, neurodegeneration, and improved cognitive function observed in our study.[34, 35]

As before, we examined the link between gene-set enrichment and concentrations of sphingolipid, but this time in the hippocampus. Results from the LC-MS analyses of hippocampal tissue was largely opposite that of the cortex (especially in the AA mice), and it shows that AA mice had high levels of the measured sphingolipids (**Supplementary Fig. 3D**). In AA+RSD mice, we observed high levels of Cer and even higher levels of SM but lower levels of HexCer, Sph, and S1P. Also, SS mice had lower levels of Cer, HexCer, SM, and S1P, except Sph compared to AA mice. We also noted that SS+RSD mice has lower levels of all sphingolipids with even lower levels of Sph. LSM remained relatively consistent across all groups. In AA+RSD+minocycline mice, we observed lower levels Cer, SM, Sph, and S1P. In SS+RSD+minocycline mice, we observed higher levels of all sphingolipids, compared to SS+RSD mice, with the exception of LSM which remained consistent across all groups.

In summary, these data support the hypothesis that sphingolipid metabolism and signaling may be involved in RSD-linked cognitive decline and neuroinflammation. However, because sphingolipid functions in the brain, depends on the brain region and may vary by fatty acid chain length, further studies are warranted. [28]

### Figure Legends

**Supplemental Fig. 1**. Hippocampal neural cell distribution in sickle and AA mice.

**Supplemental Fig. 2.** Gene set enrichment analysis showing how RSD and minocycline affect pathways, biological processes, and diseases related to cognitive function, brain development, and inflammation. SS-vs-AA_UP: genes enriched in control SS mice compared to control AA mice. SS-vs-AA_DOWN: genes downregulated in control SS mice compared to control AA mice. SS-vs-AA_Stress_UP: genes enriched in SS mice exposed to RSD compared to AA mice exposed to RSD. SS-vs-AA_Stress_DOWN: genes downregulated in SS mice exposed to stress compared to AA mice exposed to stress. SS_Drug_UP: genes enriched in SS mice treated with minocycline compared to control SS mice. SS_Drug_DOWN: genes downregulated in SS mice treated with minocycline compared to control SS mice. SS_Stress_UP: genes enriched in SS mice exposed to RSD compared to control SS mice. SS_Stress_DOWN: genes downregulated in SS mice exposed to RSD compared to control SS mice. SS_Stress_Drug_UP: genes enriched in SS mice exposed to stress and treated with minocycline compared to SS mice exposed to stress only. SS_Stress_Drug_DOWN: genes downregulated in SS mice exposed to stress and treated with minocycline compared to SS mice exposed to stress only. AA_Stress_UP: genes enriched in AA mice exposed to RSD compared to control AA mice. AA_Stress_DOWN: genes downregulated in AA mice exposed to RSD compared to control AA mice. AA_Stress_Drug_UP: genes enriched in AA mice exposed to stress and treated with minocycline compared to AA mice exposed to stress only. AA_Stress_Drug_DOWN: genes downregulated in AA mice exposed to stress and treated with minocycline compared to AA mice exposed to stress only.

**Supplemental Fig. 3.** Sphingolipids and lipid metabolism in RSD-mediated inflammation and cognitive impairment. A) Cortex GSEA results of RSD exposure and minocycline treated mice. B) Mass spectrometry characterization of sphingolipid species found in the cortex. C) Hippocampus GSEA results of RSD exposure and minocycline treated mice  D) Mass spectrometry characterization of sphingolipid species in the hippocampus
