## Supplementary figures and images for "Neuroinflammation underlies the development of social stress induced cognitive deficit in sickle cell disease"

### Supplementary Figure 1

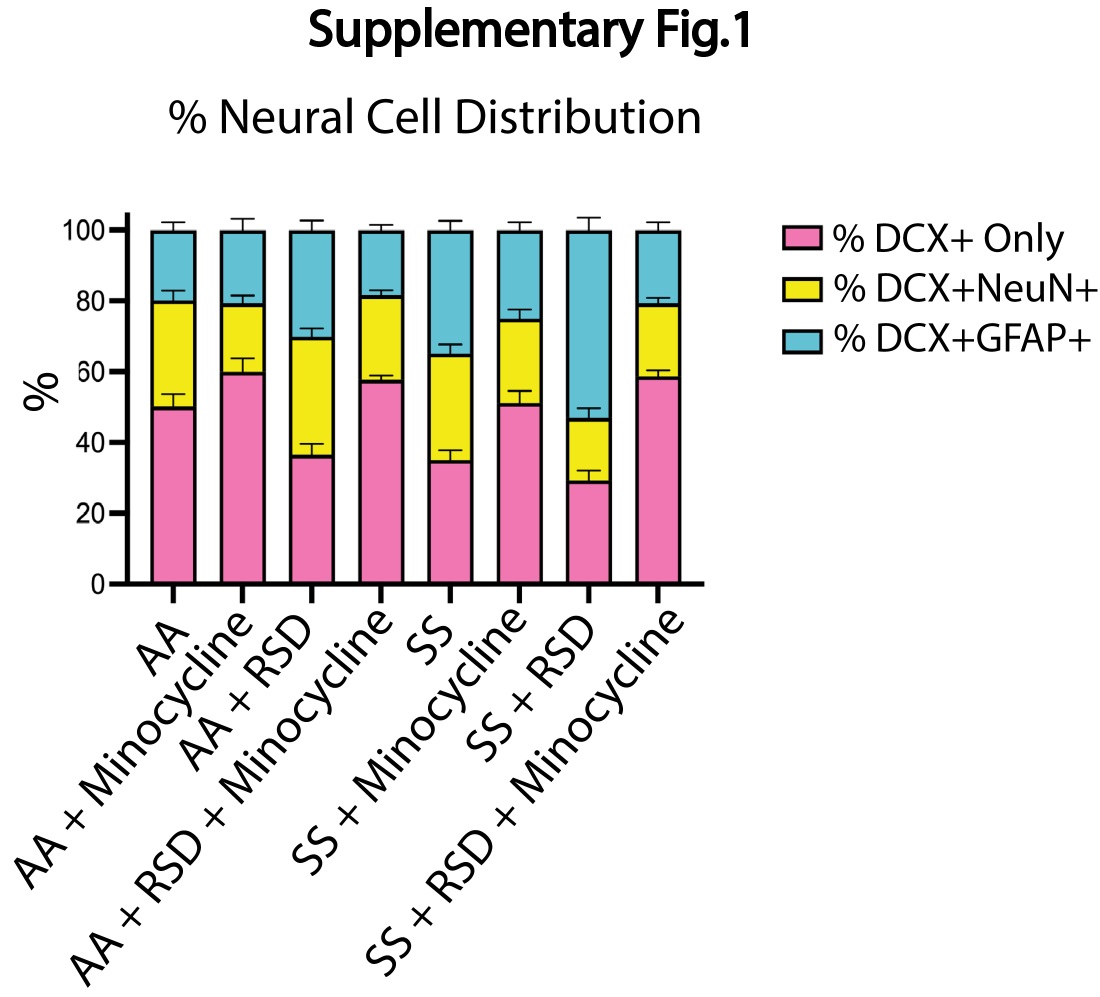

### Supplementary Figure 2

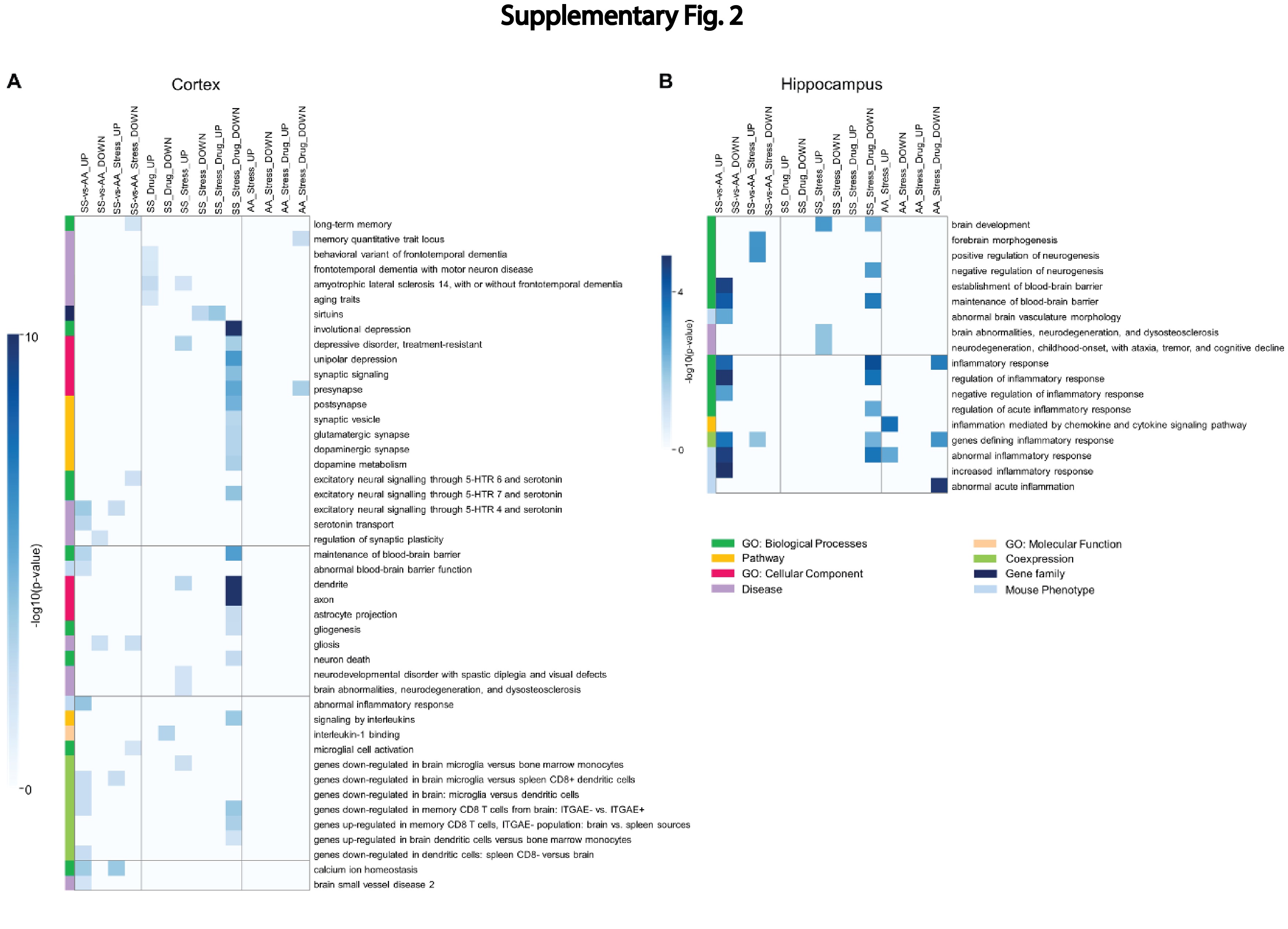

### Supplementary Figure 3

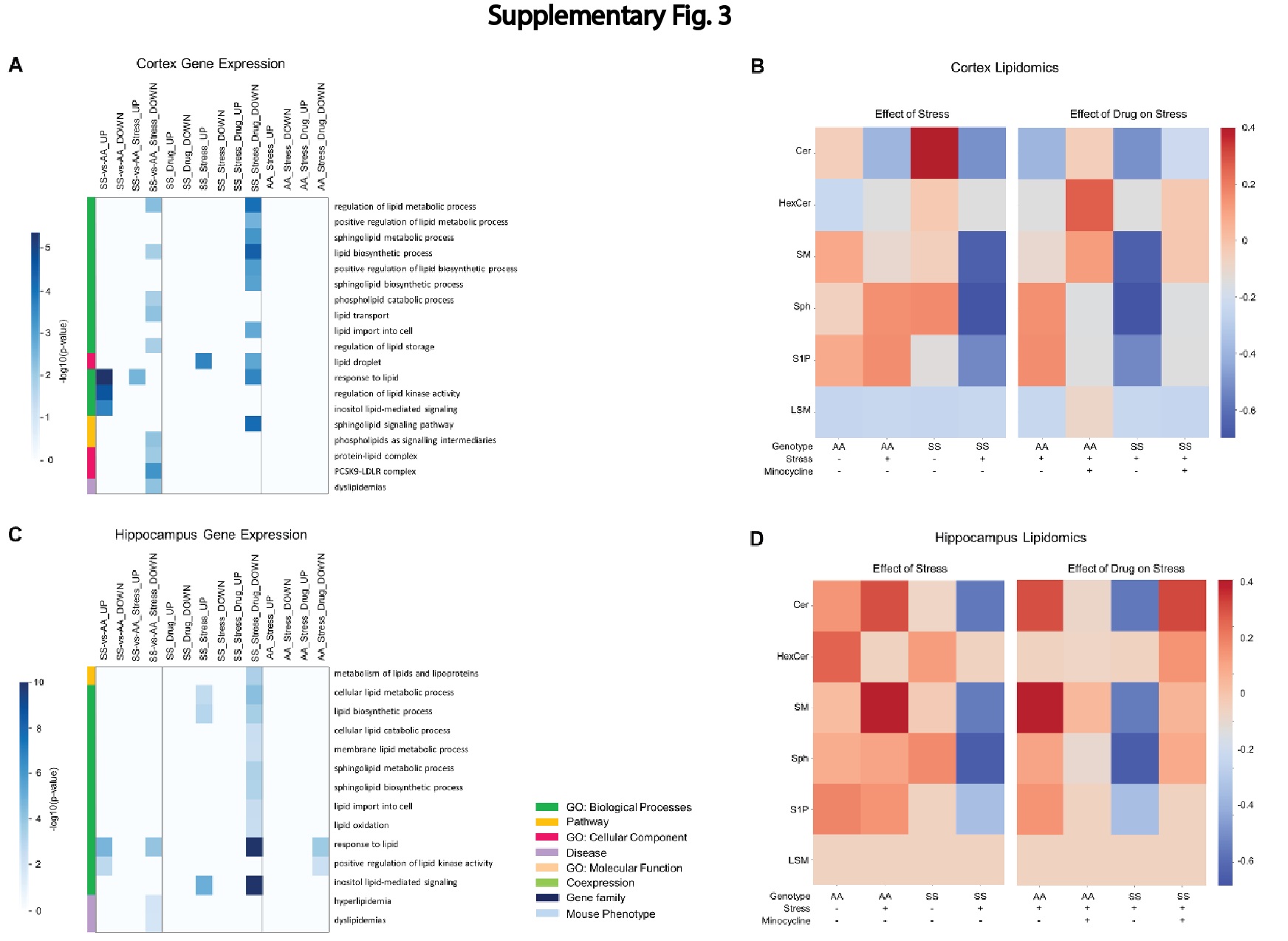
